# Alterations of oral microbiota and impact on the gut microbiome in type 1 diabetes mellitus revealed by multi-omic analysis

**DOI:** 10.1101/2022.02.13.480246

**Authors:** B.J. Kunath, O. Hickl, P. Queirós, C. Martin-Gallausiaux, L.A. Lebrun, R. Halder, C.C. Laczny, T.S.B. Schmidt, M.R. Hayward, D. Becher, A. Heintz-Buschart, C. de Beaufort, P. Bork, P. May, P. Wilmes

## Abstract

**Background:** Alterations of the gut microbiome have been linked to multiple chronic diseases. However, the drivers of such changes remain largely unknown. The oral cavity acts as a major route of exposure to exogenous factors including pathogens, and processes therein may affect the communities in the subsequent compartments of the gastrointestinal tract. Here, we perform strain-resolved, integrated multi-omic analyses of saliva and stool samples collected from eight families with multiple cases of type 1 diabetes mellitus (T1DM).

**Results:** We identified distinct oral microbiota mostly reflecting competition between streptococcal species. More specifically, we found a decreased abundance of the commensal *Streptococcus salivarius* in the oral cavity of T1DM individuals, which is linked to its apparent competition with the pathobiont *Streptococcus mutans*. The decrease in *S. salivarius* in the oral cavity was also associated with its decrease in the gut as well as higher abundances in facultative anaerobes including *Enterobacteria*. In addition, we found evidence of gut inflammation in T1DM as reflected in the expression profiles of the *Enterobacteria* as well as in the human gut proteome. Finally, we were able to follow transmitted strain-variants from the oral cavity to the gut at the metagenomic, metatranscriptomic and metaproteomic levels, highlighting not only the transfer, but also the activity of the transmitted taxa along the gastrointestinal tract.

**Conclusions:** Alterations of the oral microbiome in the context of T1DM impact the microbial communities in the lower gut, in particular through the reduction of “oral-to-gut” transfer of *Streptococcus salivarius*. Our results indicate that the observed oral-cavity-driven gut microbiome changes may contribute towards the inflammatory processes involved in T1DM. Through the integration of multi-omic analyses, we resolve strain-variant “mouth-to-gut” transfer in a disease context.

## Introduction

Thousands of distinct microbial taxa colonise the different mucosal and skin habitats of the human body^1^. These communities and their functional gene complements directly interface with host physiology, most notably the immune system^2,3^. Altered community compositions are thought to play crucial roles in triggering inflammatory processes which are most likely drivers of chronic diseases^1,4,5^, including autoimmune diseases^6–8^. The human microbiome is influenced by biotic and abiotic factors specific to each body site, which leads to distinct microbial community compositions^9^. Although closely related taxa can be present at multiple sites, most species exhibit differentiation into locally adapted strains^10^.

Bacterial species usually consist of an ensemble of strains which form coherent clades^11^. Thereby they are clearly distinguishable from the closest co-occurring related species based on their high genetic similarity^12,13^. The classical metagenomic approach consists of assembling short DNA reads into contigs and to group them into different metagenome-assembled genomes (MAGs). However, the assembly produces a patchwork of consensus contigs corresponding to the most abundant genotypes in the sample and thus can lose strain variations. Multiple approaches exist to retrieve variant information which typically involves the mapping of the metagenomic reads against the assembled contigs or reference genomes. Variant calling is then performed to determine the alleles or haplotypes^14^. Despite the genetic similarity between strains of a single species, the individual strains can exhibit different phenotypes. Such cases are notably well documented in the context of pathogenicity where many species are known to have both pathogenic and commensal strains^11^. Therefore, strain-level resolution is highly relevant in the study of the human microbiome and its links to health and disease.

The gut microbiome has been extensively studied primarily in the context of chronic diseases including cardiovascular diseases^15^, inflammatory bowel disease^16^, obesity^17^, cancers^18^, neurodegenerative diseases^19^ or autoimmune conditions such as rheumatoid arthritis^20^ or type-1^21^ and type-2 diabetes^22^. Type-1 diabetes mellitus (T1DM) is a chronic disease characterised by insulin deficiency due to autoimmune destruction of insulin-producing β-cells within the pancreatic islets. T1DM often starts during the early years of life and is one of the most common chronic diseases in childhood^23^. Its incidence worldwide has reached 15 per 100,000 people and has been globally increasing in the last decades in most developed countries^24–26^. Despite a significant genetic influence, the rise in T1DM prevalence in individuals who are not genetically predisposed strongly suggests an interplay between genetic predisposition and environmental factors^27^.

Among the possible different environmental factors, the gut microbiome modulates the function of the immune system via direct and indirect interactions with innate and adaptive immune cells^3,28^. Several studies have shown alterations of the gut microbiome composition between individuals with T1DM compared to healthy controls^29–32^. However, contrasting findings between studies have not led to a generalizable microbiome signature for T1DM and it still remains unclear how microbiome changes affect the gastrointestinal tract and immune functions in T1DM.

The oral cavity and the colon sit at opposite sides of the gastrointestinal tract. The mouth is considered a gateway to different organs of the body, and therefore acts as a potential reservoir for different pathogens^33^. Poor dental health and dysfunctional periodontal immune-inflammatory reactions caused by bacterial pathogens may lead to periodontitis and are associated with increased risks of developing systemic inflammatory disorders^34^. The development of inflammation in the oral cavity has notably been found to be associated with systemic inflammation and cardiovascular disease^35^, insulin resistance^36^ and complications in type-1 and type-2 diabetes^37^. Despite the limited number of shared taxa between the oral cavity and the lower gut^38^ due to the gastric bactericidal barrier, intestinal motility or bile and pancreatic secretions^39^, a recent study has shown that the oral community type was predictive of the community recovered from stool^40^. Additionally, Schmidt, Hayward *et al*. recently found that a subset of 74 species were frequently transmitted from mouth to gut and formed coherent strain populations along the gastrointestinal tract^41^. Finally, it is known that the physiology of the oral cavity is altered in T1DM patients, notably with a decrease of salivary flow rate (dry-mouth symptom) and an increased concentration of glucose in the saliva and subsequent acidification of the oral cavity^42–44^. However, the effect of T1DM on the microbiome of the oral cavity, or the effect of the microbiome on T1DM in general is still poorly understood, with few, and regularly contradicting findings^45^.

Here, we apply an integrated multi-omic approach to characterise differences in the oral and gut microbiomes in the context of T1DM. We identify distinct oral microbiota suggestive of competition between streptococcal species and an acidified oral cavity. We link these differences to alterations in the gut microbiome and the host’s inflammatory response. Finally, we explore the level of mouth-to-gut transmissions in T1DM, highlight transferred and active strains, and identify differences in strain-level transmission profiles in T1DM patients compared to healthy controls.

## Methods

### Ethics

Written informed consent was obtained from all subjects enrolled in the study. This study was approved by the Comité d’Ethique de Recherche (CNER; reference no. 201110/05) and the National Commission for Data Protection in Luxembourg.

### Sample acquisition

The study design was an observational study of eight selected families (M01-M06, M08, M11) containing at least two members with T1DM and healthy individuals in two generations or more, from existing patient cohorts from the Centre Hospitalier du Luxembourg. Individual patients are annotated as a combination of their family and a number for each individual per family (*e*.*g*. M05.1). Recruited families were seen three times (V1, V2, V3) at intervals of between four and eight weeks for data and samples collection. Donors collected 2–3 ml of saliva at home before dental hygiene and breakfast in the early morning. Faecal samples were also self-collected and both samples were immediately frozen on dry-ice, transported to the laboratory and stored at -80°C until further processing. Part of the cohort’s raw data (families M01-04)^41,46^ as well as the oral and gut metagenomics (families M05-11)^41,46^ were previously studied and published. The following method sections describe the processing of the newly produced dataset.

### Biomolecular extractions

For each individual and visit, faecal and saliva samples were subjected to comprehensive biomolecular isolations.

For the faecal samples, 150 mg of each snap-frozen sample was reduced to a fine powder and homogenised in a liquid nitrogen bath followed by the addition of 1.5 ml of cold RNAlater and brief vortexing prior to incubation overnight at -20°C. After incubation, the sample was re-homogenised by shaking for 2 minutes at 10 Hz in an oscillating Mill MM 400 (Retsch) and subsequently centrifuged at 700 *x* g for 2 minutes at 4°C. The supernatant was retrieved and the cells were pelleted by centrifugation at 14,000 *x* g for 5 minutes. Cold stainless steel milling balls and 600 µl of RLT buffer (Qiagen) were added to the pellet and this was re-suspended via quick vortexing. Cells were disrupted by bead beating in an Oscillating Mill MM 400 (Retsch) for 30 seconds at 25 Hz and at 4°C. Finally, the lysate was transferred onto a QIAshredder column and centrifuged at 14,000 *x* g for 2 minutes and the eluate retrieved for multi-omics extraction. The subsequent biomacromolecular extractions were based on the Qiagen Allprep kit (Qiagen) using an automated robotic liquid handling system (Freedom Evo, Tecan) as described in Roume *et. al* and in accordance with the manufacturer’s instructions^47^. For the saliva samples, the individual snap-frozen sample was thawed on ice, and 1 ml was subsampled and centrifuged at 18,000 *x* g for 15 minutes at 4°C. The supernatant was discarded and the pellet directly refrozen in liquid nitrogen. Cold stainless steel milling balls were added to the frozen pellet for homogenisation by cryo-milling for 2 minutes at 25 Hz in an oscillating Mill MM 400 (Retsch). Subsequently, 300 µl of methanol and 300 µl of chloroform were added before a second passage through the Oscillating Mill at 20 Hz for 2 minutes. After centrifugation at 14,000 *x* g for 5 minutes, two phases (polar and non-polar) and a solid interphase were visible. The two phases were discarded and the solid interphase kept for multi-omics extraction. Stainless steel milling balls and 600 µl of RLT buffer (Qiagen) were added to the pellet, re-suspended via quick vortexing and cells were disrupted by bead beating in an Oscillating Mill MM 400 (Retsch) for 30 seconds at 25 Hz at 4°C. The lysate was transferred onto a QIAshredder column and centrifuged at 14,000 *× g* for 2 minutes. The subsequent steps were performed as described for the faecal samples.

### DNA sequencing

After extraction, the retrieved DNA was depleted of leftover RNA by RNAse A treatment at 65°C for 45 minutes. After ethanol precipitation, the samples were re-suspended in 50 µl nuclease-free water. The quality and quantity of the retrieved DNA were assessed both before and after treatment via gel electrophoresis and Nanodrop analysis (ThermoFisher Scientific). Sequencing libraries for salivary samples were prepared using the NEBNext Ultra DNA Library Prep kit (New England Biolabs, Ipswich) using a dual barcoding system, and sequenced at 150 bp paired-end on Illumina HiSeq 4000 and Illumina NextSeq 500 machines.

### RNA sequencing

The extracted RNA was treated with DNase I at 37°C for 30 minutes and purified using phenol-chloroform. From the aqueous phase, RNA was precipitated with isopropanol and re-suspended in 50 µl nuclease free water.

RNA integrity and quantity were assessed before and after treatment using the RNA LabChip GX II (Perkin Elmer). Subsequently, 1 µg of RNA sample was rRNA-depleted using the RiboZero kit (Illumina, MRZB12424). Further library preparation of rRNA-depleted samples was performed using TruSeq Stranded mRNA library preparation kit (Illumina, RS-122-2101) according to the manufacturer’s instructions apart from omitting the initial steps for mRNA pull-down. Prepared libraries were checked again using the RNA LabChip GX II (Perkin Elmer) and quantified using Qubit (Invitrogen). A 10 nM pool of the libraries was sent to the EMBL genomics platform for sequencing on a Illumina NextSeq 500 machine.

### Protein processing and mass spectrometry

The following section describes the procedures for samples from families M05, M06, M08, and M11. For a description of the protein processing of samples from families M01-M04 see Heintz-Buschart *et al*. 2016^46^.

Extracted proteins were processed and digested using the S-Trap™ system (ProtiFi) following manufacturer’s instructions. Briefly, protein suspensions were solubilised with SDS then reduced, alkylated and acidified for complete denaturation.

Approximately 200 μl of samples were transferred onto the S-Trap column and centrifuged until all of the sample volume was transferred. The columns were then washed twice with 180 μl S-Trap protein binding buffer. Protein digestion was performed by adding 20 μl of 0.04 μg/μl trypsin solution to each column, to achieve a trypsin to protein ratio of 1:50. Incubation was performed for three hours at 47°C in a Thermomixer. Tryptic peptides were eluted with 40 μl 50 mM TEAB, 40 μl 0.1% acetic acid, and 35 μl 60% acetonitrile with 0.1% acetic acid at 4,000 *x g* for 1 minute per elution. Samples were dried at 45°C in a vacuum centrifuge and stored at -20°C.

Peptides were fractionated into eight fractions using the high pH reversed-phase peptide fractionation kit (Pierce™ Thermo Fisher Scientific) according to the manufacturer’s instructions and using self-made columns as previously described^48^. Digested, dried peptides were resuspended in 300 μl of 0.1% trifluoroacetic acid and suspensions transferred onto the columns. After centrifugation at 3,000 *x g* for 2 minutes the eluate was retained as “flow-through”-fraction. Columns were then washed with 300 μl water (ASTM Type I) at 3,000 *x g* for 4 minutes. Separation of samples into eight fractions was performed using 300 μl of elution solutions with increasing concentrations of acetonitrile in 0.1% trifluoroacetic acid at 3,000 *x g* for 4 minutes. Each elution fraction was collected in a separate microcentrifuge tube, dried at 45°C in a vacuum centrifuge and stored at -20°C.

Peptide concentrations were measured for fraction two of each sample using the Quantitative Fluorometric Peptide Assay kit (Pierce™ Thermo Fisher Scientific) according to the manufacturer’s instructions.

Of each of the samples, for each fraction, the volume for 170 ng of peptides were loaded onto in-house built columns (100 μm x 20 cm), filled with 3 μm ReproSil-Pur material and separated using a non-linear 100 min gradient from 1 to 99% buffer B (99.9% acetonitrile, 0.1% acetic acid in water (ASTM Type I) at a flow rate of 300 nl/min operated on an EASY-nLC 1200. Measurements were performed on an Orbitrap Elite mass spectrometer performing one full MS scan in a range from 300 to 1,700 m/z followed by a data-dependent MS/MS scan of the 20 most intense ions, a dynamic exclusion repeat count of 1, and repeat exclusion duration of 30s.

### Metagenomic and metatranscriptomic data analysis

For each individual time point, metagenomic (MG) and metatranscriptomic (MT) data were processed and co-assembled using the Integrated Meta-omic Pipeline (IMP)^49^ which includes steps for the trimming and quality filtering of the reads, the filtering of rRNA from the MT data, and the removal of human reads after mapping against the human genome (hg38). Pre-processed DNA and RNA reads were co-assembled using the IMP-based iterative co-assembly using MEGAHIT 1.0.3^50^. After co-assembly, prediction and annotation of open-reading frames (ORFs) were performed using IMP and followed by binning and then taxonomic annotation at both the contig and bin level. MG and MT read counts for the predicted genes obtained using featureCounts^51^ were linked to the different annotation sources (KEGG^52^, Pfam^53^, Resfams^54^, dbCAN^55^, Cas^56^, and DEG^57^, as well as to taxonomy (mOTUs 2.5.1^58^ and Kraken2 using the maxikraken2_1903_140GB database^59^). Kraken2 annotations were used to generate read count matrices for each taxonomic rank (Phylum, Class, Order, Family, Genus and Species) by summing up reads at the respective levels.

### Identification of variants

IMP produced the mapping of the processed DNA and RNA reads against the final co-assembled contigs with the Burrows-Wheeler Aligner tool (BWA 0.7.17)^60^ using the BWA-MEM algorithm with default parameters. Additionally for each individual, the oral DNA reads from all available visits were mapped against the gut contigs produced from all available visits with the same parameters.

All alignment files per sample were used to call variants using bcftools 1.9^61,62^. Bcftools *mpileup* was run on the gut contigs as reference FASTA file with default parameters except for the *--max-depth* being set to 1000 to increase variant calling certainty. Called variants were filtered based on their quality and read depth with minimum values set to 20 and 10, respectively and indels were excluded. Subsequently, in order to reinforce confidence in the variant calling, variants were kept for downstream analysis, only if they fitted the following criteria: (i) positive allelic depths on both the forward and reverse strands for the corresponding gut and oral DNA reads, and (ii) presence of an alternative allele (genotype=1 in the vcf file) at the oral DNA reads and the gut RNA read levels. These criteria ensured that the variants were resolved in both the gut and oral samples at both the DNA and RNA levels.

After variant identification and filtering, reads containing the variants were extracted from the mapping files and taxonomically annotated using Kraken2 for sample comparison. For metaproteomics, missense variants (variant that leads to a different amino acid) were identified using a previous in-house script^46^ and the generated ORFs containing variants were added to the metaproteomic database (see below).

### Metaproteomic data analysis

As the mass spectrometry analysis of the protein fraction was performed at different facilities for families M01-04 and families M05, M06, M08, and M11, certain parts of the preprocessing workflow and analyses had to be tailored to the data, as mentioned below.

Raw files were converted to mzML format using ThermoRawFileParser^63^ and to ms2 format using ProteoWizard’s msconvert^64^. The files for families M01-04 were filtered for the top 300 most intense spectra, the files for the other families for the top 150 most intense spectra to optimise protein identifications.

For each sample, microbial protein sequence databases were constructed from the Prokka^65^ predicted protein sequences of the IMP co-assemblies and supplemented with variant protein sequences (missense variants) identified in both the oral cavity and the gut, during the variant calling step. This was done in order to consider only the variant sequences originating from the oral cavity that could also be found in the gut. If no database was available for a single sample, all databases available from the individual were concatenated. If an individual had no database, all databases from the individual’s family were concatenated. In addition, the human RefSeq protein sequences (release 92), a collection of plant storage proteins that might be present due to food intake as well as the cRAP contaminant database (release 04/03/2019) were added. The databases were then filtered according to size (60-40000 residues) to eliminate noise from very large or small proteins that can be erroneously produced during the ORF prediction step. Duplicate sequences were removed by sequence using SeqKit^66^.

Concatenated target-decoy databases were built using Sipros Ensembles *sipros_prepare_protein_database*.*py*. Using Sipros Ensemble^67^, each sample was searched against the prepared database for that sample. Identifications were filtered to a protein FDR of 1%.

After the search, human and microbial protein identifications were treated separately. Human proteins/protein groups that ended up having identical protein identifiers after processing the database identifiers in the output were collapsed and their spectral counts summed up. The same was done for the microbial proteins but gene identifiers were replaced by the corresponding annotation identifiers from the respective source (e.g., KEGG, Pfam, see above).

### Statistical analyses

An initial screening was performed based on MG and MT sequencing and assembly statistics, principal component analysis and hierarchical clustering on gene abundances to highlight potential outliers. Samples whose sequencing and assembly statistics consistently appeared outside +/-1.5x the interquartile range and clustered substantially differently compared to other samples from the same individual with hierarchical clustering were considered as outliers and removed from the dataset. Similarly, filtering was performed for the MP data with MS raw data quality and protein identification rate. After quality control, several individuals were removed because of their high variability due to either a very young age (age under 4 years old for M08-04 and M11-03) or a comorbidity that was not present in the rest of the dataset (T2DM for M11-05 and M11-06).

After taxonomic and functional analysis, gene/taxa read count and protein spectral count matrices were generated for differential abundance and expression analysis using the DESeq2 R package^68^. As the sampling visits for each individual are not independent, the median value for each gene/protein of the available visits for each individual was computed to obtain a matrix with one representative value per gene/protein per individual. Additionally, genes in read count matrices were removed if they did not have at least 20 reads in 25% of all the individuals, ensuring sufficient representation of the gene in the sample set for downstream statistical analyses. Proteins in the spectral count matrices were removed if they did not have at least ten spectra in 25% of all the individuals. Finally, family membership was set as confounder for the DESeq2 the differential analyses.

For the correlation analyses, Spearman’s rank correlation coefficients were calculated with two-sided significance tests, corrected for multiple testing using the Benjamini-Hochberg method with a p-value of 0.001 and a significance threshold of 0.7, using the *rstatix* R package (https://github.com/kassambara/rstatix).

### Diversity analysis

Raw counts were transformed from absolute counts to relative abundances by dividing each value by sample total abundance. The richness as a total number of detected species after filtering was recorded as well as alpha diversity using the Simpson index^69^. Beta diversity was analysed using Bray-Curtis as a distance measure with hierarchical clustering, distance-based redundancy analysis (dbRDA), and nonmetric multidimensional scaling (NMDS). Significance tests between groups were carried out using the Mann–Whitney–Wilcoxon test (MWW) or analysis of variance (ANOVA, dbRDA formula: *species ∼ condition+family*). Analyses were performed in R using the *picante*^*70*^ and *vegan*^*71*^ packages.

## Results and Discussions

### Study description

In this study, we performed a multi-omic oral and gut microbiome study of eight families with at least two T1DM cases per family (**Figure 1A**). This expanded on previous studies focussing on a subset of the data^41,46^. The present work additionally includes metagenomic (MG) and metatranscriptomic (MT) analyses of the oral cavity for all participants. In total, we analysed 84 stool and 76 saliva samples from 35 individuals. We generated MG, MT and MP data for each individual from multiple visits. Of the 35 individuals, 17 were T1DM patients and 18 were healthy family members (**Figure 1A**). In total, 653.4 Giga base pairs (Gbp) of DNA sequencing data, 870.6 Gbps RNA sequencing data, and 13,833,325 fragment ion spectra were acquired.

**Figure 1.**
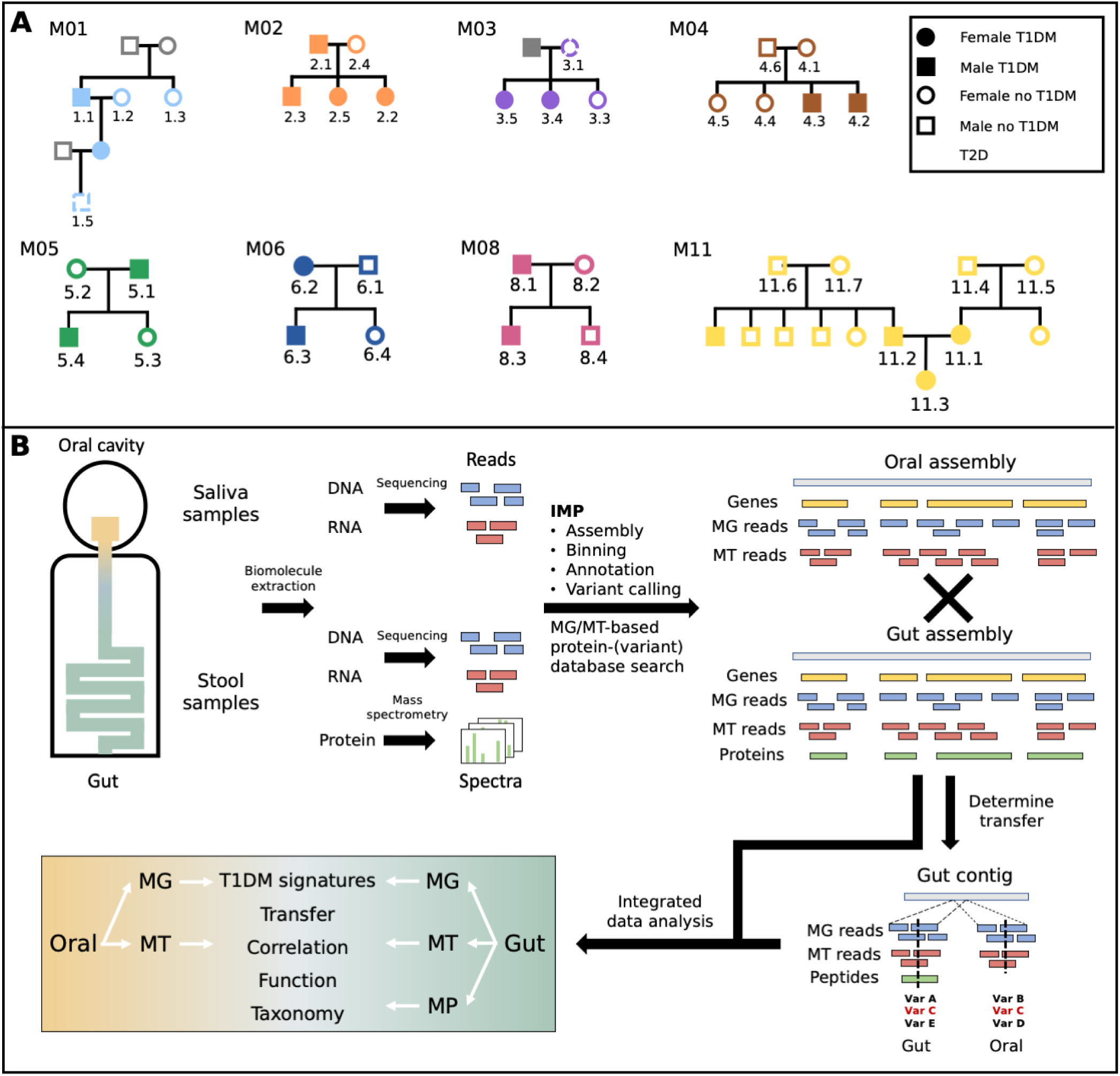
Description of the cohort and overview of the study workflow. The upper panel(**A**) shows the different individuals with family membership as well as disease status in the cohort. The lower panel (**B**) describes the integrated multi-omics analysis workflow to process, integrate and analyse metagenomic (MG), metatranscriptomic (MT) and metaproteomic (MP) data from saliva and stool samples.

**Table 1.** Overview of the multi-omics study data. Number of samples used per ome, average values of sequencing and mass spectrometry (MS) raw data and key metrics from saliva and stool samples. Sequencing data acquired are shown in Giga base pairs (Gbp), percentage of recruitment of sequencing reads to assemblies, fragment ion spectra acquired (MS2 scans), and protein groups identified. All values are shown with standard deviation. MG: Metagenomics, MT: Metatranscriptomics, MP: Metaproteomics, k: kilo.

|  | MG samples | MG sequencing data [Gbp] | MG read recruitment [%] | MT samples | MT sequencing data [Gbp] | MT read recruitment [%] | MP samples | MP MS2 scans [k] | MP protein groups [k] |
| --- | --- | --- | --- | --- | --- | --- | --- | --- | --- |
| **Stool** | 84 | 4.2 ± 0.8 | 95.3 ± 2.5 | 64 | 5.6 ± 1.1 | 56.8 ± 9.8 | 71 | 94.7 ± 59.4 | 4.5 ± 3.5 |
| **Oral** | 74 | 4.2 ± 0.9 | 83.5 ± 12.1 | 74 | 7.0 ± 1.6 | 62.7 ± 16.7 | - | - | - |

Over all samples, the DNA and RNA sequencing data per sample amounted to on average 4.2 ± 0.9 Gbp for MG and 6.3 ± 1.6 Gbp for MT. While the gut data consisted of 4.2 ± 0.8 Gbp of MG and 5.6 ± 1.1 Gbp MT sequencing data, the oral data represented 4.2 ± 0.9 Gbp of MG and 7.0 ± 1.6 Gbp MT sequencing data. For the stool samples, on average 95,000 ± 59,000 MS2 scans were performed and 4,500 ± 3,400 proteins identified. For samples from families 01-4 on average 63,000 ± 4,700 fragment ion scans were obtained. The database searches resulted in 1,500 ± 300 proteins on average. A mean of 203,000 ± 11,800 fragment ion scans were obtained for samples from families 05, 06, 08, and 11 and 8,000 ± 1,600 proteins could be identified. For detailed statistics see **Supplementary Table 1**. In the present study, we combined information from three omes in order to identify and follow strain-variants across the two body sites. To be able to do so, the overlap among the different omes had to be maximised to preserve all their sample specificity. Thus, the complete set of contigs from sample-specific assemblies were used rather than metagenome-assembled genomes that would have only covered a subset of all the multi-omic data (**Figure 1B**).

### Overall microbial community structure does not differ significantly between T1DM and healthy controls

We compared the community structures of both body sites between T1DM patients and controls using the MG data. Overall, the number of total species detected in the gut varied more in healthy individuals, but no significant differences in richness (MWW: p-val 0.72, **Supplementary Figure 1A**) nor in Simpson’s Index of Diversity were observed (MWW: p-val 0.53, **Supplementary Figure 1B**). Beta diversity differed significantly according to family membership but not between T1DM patients and controls (ANOVA on dbRDA; p-vals: 0.001 (family), 0.11 (condition); R^2^: 0.49; **Supplementary Figure 1C**).

The oral microbiota did not differ significantly in species richness (MWW: p-val 0.48, **Supplementary Figure 1A**) nor in their Simpson’s Index of Diversity (MWW: p-val 0.90, **Supplementary Figure 1B**). The beta diversity, as in the gut, showed no significant difference for T1DM but for family membership (ANOVA on dbRDA, p-vals: 0.5 (condition), 0.003 (family); R^2^: 0.37; **Supplementary Figure 1C**). Thereby, for both body sites, no evidence was found that suggested a significant effect of T1DM on the overall microbiota community diversity. As shown before, observable differences in oral community composition may instead be related to family membership^46^.

### The acidification of the oral cavity in T1DM impacts specific taxa and destabilises the equilibrium between Streptococcus species

*Streptococcus* species are the primary colonisers of the oral cavity and are key players in oral homeostasis and disease^72^. In healthy subjects, there is a balance between the abundance of opportunistic pathogens (*e*.*g. S. mutans* or *S. pneumoniae*) and non-pathogenic commensal species (*e*.*g. S. salivarius, S. parasanguinis* or *S. mitis)* which compete with each other via different mechanisms such as acid or base production, or secretion of bacteriocins^72–75^.

In our study, the abundance of several members of the genus *Streptococcus* varied in the oral cavity of T1DM patients compared to controls. In particular at the MG level, we observed high variability among *Streptococcus* species (**Figure 2**). Such variability is in agreement with previous findings whereby the numbers of different *Streptococcus* species were found to be increased or decreased in T1DM depending on the study^76,77^. For example, a 16S rRNA gene-based study of both body sites observed an increase in the abundance of the genus *Streptococcus* in the mouth but a decrease in the gut of T1DM patients^45^.

**Figure 2.**
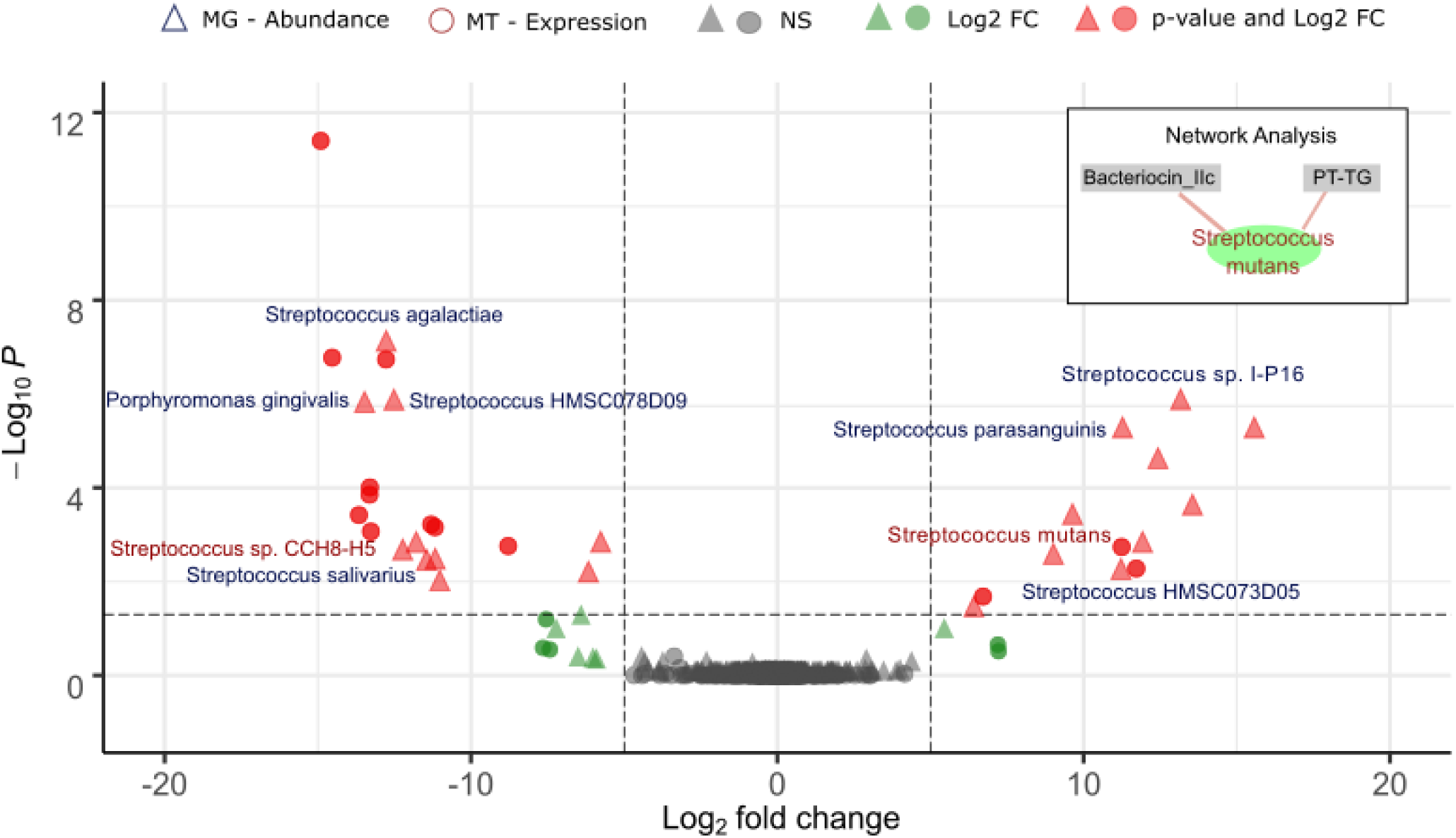
Taxon-resolved differential abundance and gene expression in the oral microbiome in T1DM. The differences in abundance (triangles) and expression (circle) in T1DM versus healthy individuals using metagenomic and metatranscriptomic data, respectively, are shown on the volcano plot. A minimum log_2_ fold change of 5 (dashed vertical lines) and an adjusted p-value of 0.01 (dashed horizontal line) were required (red dots). Taxa that satisfy the fold-change threshold but not the adjusted p-value threshold are displayed in green. A subset of **Supplementary Figure 2** is shown in the insert in the upper-right and highlights the correlation between *S. mutans* activity and the expression of a target-specific bacteriocin.

We observed an increased abundance of the acid-tolerant but non-pathogenic *Streptococcus parasanguinis* and closely related *Streptococcus* HMSC073D05 (log_2_ fold changes 3.5 and 3.4, respectively; adj. p-val<0.05). In contrast, the abundance of the commensal and acid-intolerant *Streptococcus salivarius* was found to be decreased in T1DM (log_2_ fold change -3.5; adj. p-val<0.05)^78^. Additionally, we observed a decreased abundance of *Porphyromonas gingivalis* in the cavity of T1DM patients. *P. gingivalis* is usually associated with a dysbiotic state but is also known to be unable to grow in acidic conditions^79^. Taken together, these results indicate a microbial profile corresponding to an acidified cavity in the case of T1DM patients^42–44,80^.

Further evidence was provided by the metatranscriptomic data, which showed a significantly increased activity of the pathogenic *Streptococcus mutans*^*81*^ (log_2_ fold change 11.3 ; adj. p-val<0.05), while other *Streptococci*, notably *S. salivarius/S*. sp. CCH8-H5 (log_2_ fold change - 13.3 at adj. p-val<0.05) were less active (**Figure 2**). *S. mutans* is a common pathogen of the oral cavity associated with periodontal diseases and known for its acid-tolerance and acidogenicity, which leads to further microbial acidification of the oral cavity in T1D patients^82,83^.

In order to better understand the underlying patterns in the oral microbiomes, we looked at correlations of the expressed genes with the taxa that were found to be differentially active. We observed significant positive correlations (rho>0.7 at p-value <0.001) between *S. mutans* and two specific expressed transcripts related to bacterial competition among closely related species, namely bacteriocin IIc and pre-toxin TG, which are the constituent domains of uberolysin (**Figure 2 - Network Analysis** and **Supplementary Figure 2**). This peptidic toxin is a circular bacteriocin characterised in the genus *Streptococcus* and has a broad spectrum of inhibitory activity, which includes most streptococci with the notable exception of *S. rattus* and *S. mutans*^*84,85*^. The corresponding gene expression was not found to be linked with a particular species. However, the fact that *S. mutans* is resistant to the toxin and the observation that *S. mutans* is strongly correlated with both transcripts for this toxin, supports our hypothesis that *S. mutans* is responsible for the expression of the bacteriocin. The acidified oral cavity of T1DM patients, originally due to the host pathophysiology^42–44^, according to our data, leads to the decreased abundance of acid-intolerant bacteria and favours the growth of acid-tolerant pathogenic *S. mutans*, which then further acidifies the environment and outcompetes the commensal *S. salivarius* by expressing a target-specific bacteriocin.

### Streptococcus salivarius’ abundance decreases in the gut favouring an inflamed environment and an enterobacterial bloom

The differential abundance analysis of the gut-derived multi-omic data showed few differences between conditions. The lower abundance of *S. salivarius* in the gut follows the trend we observed in the oral cavity (**Supplementary Table 2**). *S. salivarius* colonises the intestine of adults and contributes to gut homeostasis by anti-inflammatory effects as well as by preventing the bloom of pathogens^86–88^. Previous studies have shown that a *S. salivarius* strain isolated from the oral cavity was able to prevent inflammatory responses both *in vitro* and *in vivo* by significantly reducing the activation of NF-κB and IL-8 secretion in intestinal epithelial and immune cell lines^86,89,90^. Therefore, a decrease of *S. salivarius* abundance may culminate in a more inflamed gut environment.

We also observed an increased abundance in the *Escherichia coli* (*Enterobacteria*) in the gut (**Supplementary Table 2**). *Enterobacteria* are among the most commonly overgrowing potential pathobionts whose expansion is associated with many diseases and, in particular, inflammation^91^.

By investigating gene expression in the gut, we found multiple differentially expressed genes in T1DM in comparison to healthy controls (**Figure 3**). Strikingly, a majority of the overexpressed genes are associated with *Enterobacteria* indicating a strong activity of this group in T1DM patients. They are usually found in low abundance in the gut in close proximity to the mucosal epithelium due to their facultative anaerobic metabolism^92^. *Enterobacteria* are also well known to have their growth favoured in many conditions involving inflammation^93^. The identified overexpressed genes contribute to bacterial virulence, oxidative stress response, cell motility and biofilm formation, and general replication and growth. Notably, an upregulation of a catalase-peroxidase was identified, an enzyme that detoxifies reactive oxygen intermediates such as H_2_O_2_ and, thus, is involved in protection against oxidative stress produced by the host. Enzymes associated with biofilm formation (YliH) were also overexpressed. Finally, OmpA-like transmembrane domain was identified as well the protein HokC/D, which corresponds to the *E. coli* toxin-antitoxin system that ensures the transmission of the associated plasmid.

**Figure 3.**
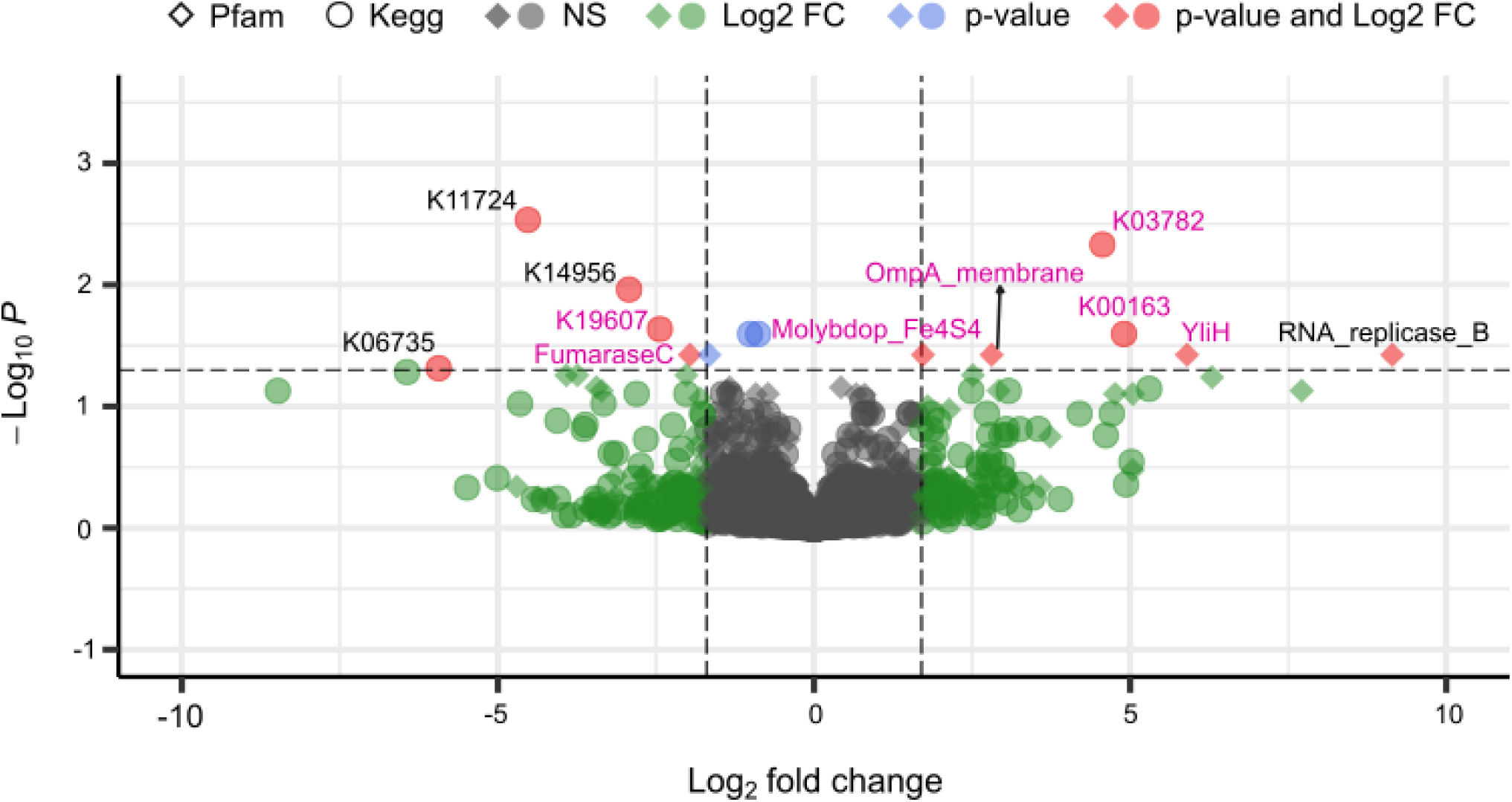
Differential gene expression analysis within the gut in T1DM. Difference in expression using metatranscriptomic data is shown on the volcano plot. A minimum log_2_ fold change of 2.5 (dashed vertical lines) and adjusted p-value of 0.05 (dashed horizontal line) were required (red dot). Functions that satisfy only the fold change or the adjusted p-value threshold are displayed in green and blue, respectively. Diamonds and circles respectively indicate complementary annotations from both the Pfam and KEGG databases. Genes associated with Enterobacteria are marked in pink.

There are multiple possible mechanisms of inflammation-driven blooms of *Enterobacteria* in the gut. One of them relies on the inflammatory host response that produces a potent antimicrobial agent (peroxynitrite) which is quickly converted to nitrate and can then be used for bacterial growth through nitrate respiration^93^. Since the genes encoding nitrate reductase in the gut are mostly encoded by *Enterobacteria*, this nitrate-rich environment provides a growth advantage for *Enterobacteria* such as *E. coli*. In addition to the genes involved in oxidative stress, we also found the molybdopterin oxidoreductase 4Fe-4S domain to be overexpressed in T1DM (**Figure 3** and **Supplementary table 3**). This domain is found in a number of reductase/dehydrogenase families and notably the respiratory nitrate reductase in *E. coli* which further supports our hypothesis of inflamed gut in the context of T1DM. Increased abundance of *Enterobacteria* in T1DM has been partially observed before but the signal was not necessarily clear^94^ or was associated with confounding factors like antibiotic-induced acceleration of T1DM^95^ and no functional evidence were found.

Additionally, we looked at the effect of T1DM on the abundance of human proteins in the gut. We hypothesised that inflammation of the gut would lead to higher abundances of proteins involved in the host immune response. Interestingly, we mostly found evidence of exocrine pancreatic insufficiency with several types of proteases, such as pancreatic carboxypeptidases, elastases or trypsin-related enzymes, being less abundant in T1DM (**Figure 4** and **Supplementary Table 4**) which can be associated with T1DM^96^. One protein involved in the host immune response, the polymeric immunoglobulin receptor (pIgR), was found at elevated levels in T1DM (log_2_ fold change 0.42 at p-val<0.05) (**Figure 4** and **Supplementary Table 4**). pIgR is a transmembrane protein expressed by epithelial cells and responsible for the transcytosis of the secreted polymeric IgA produced in the mucosa by plasma cells to the gut lumen^97,98^. Binding of polymeric IgA to the microbial surface protects the intestinal mucosa by preventing attachment to the epithelial cells, thus inhibiting infection and colonisation. When looking at differentially expressed proteins taking all individuals and visits into account, we found similar proteins as when using median information but also several additional proteins associated with the host immune response and inflammation to be more expressed in T1DM (**Supplementary Figure 3** and **Supplementary Table 5**). While that approach is statically less robust (see methods) it allows to observe additional trends in the dataset. Notably, we found higher levels of the lipocalin 2 enzyme (LCN2) (log_2_ fold change 0.37 at p-val<0.05) which is a typical biomarker in human inflammatory disease^99^ and has been associated with metabolic disorders such as obesity and diabetes^100–102^. The analysis also confirmed the higher expression of the lactotransferrin (LTF) (log_2_ fold change 0.76 at p-val<0.05), which was already found in our previous study^46^. LTF plays a role in innate immunity and insulin function^103,104^ and its antimicrobial activity can influence the gastrointestinal microbiota^105^.

**Figure 4.**
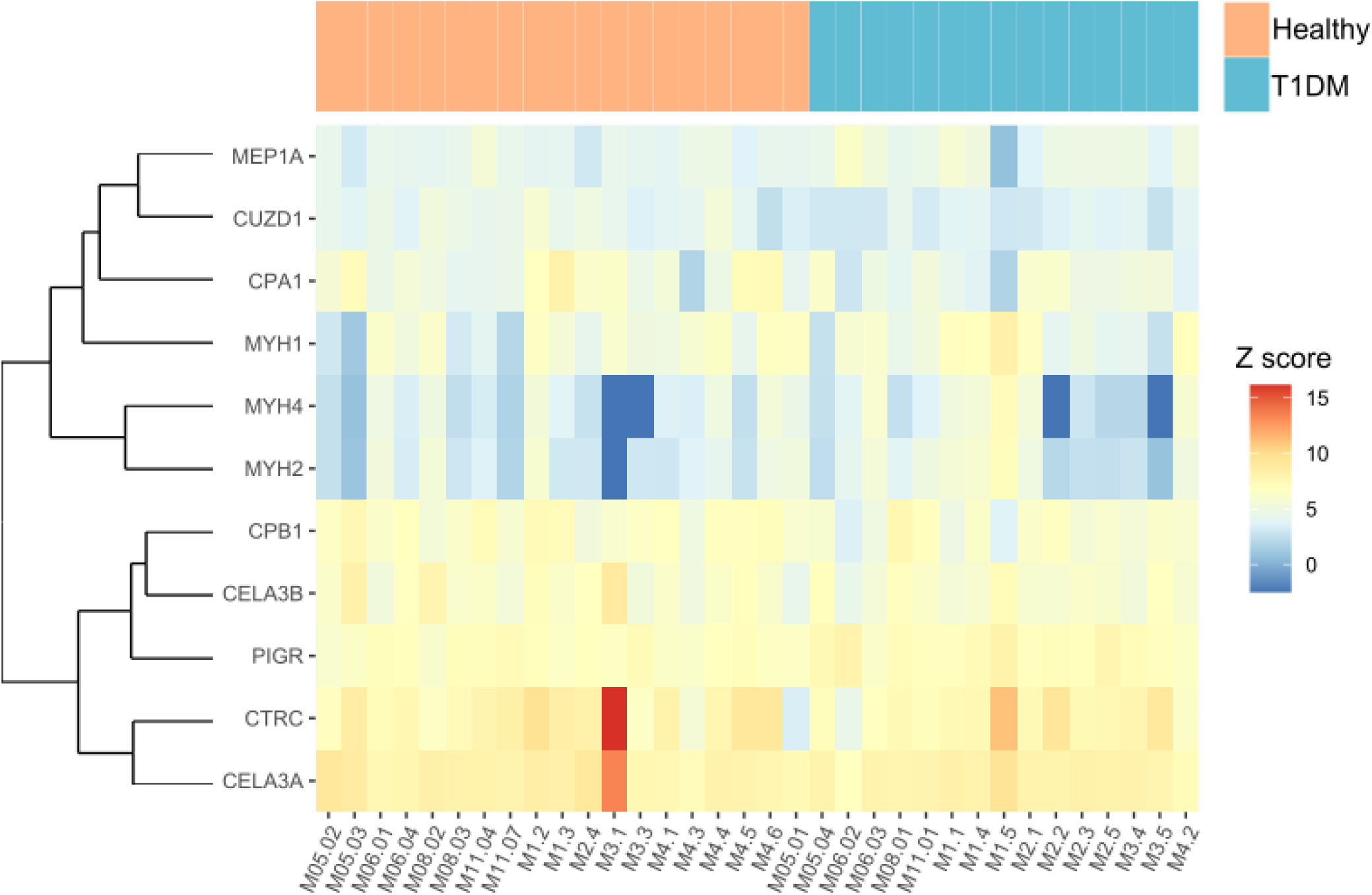
Human proteome differences in T1DM. Heatmap displaying the relative abundances of human proteins with the highest significance in a differential analysis of T1DM versus healthy individuals (unadjusted p-value < 0.05). The samples are ordered by conditions. Healthy individuals and T1DM patients are respectively shown in orange and blue boxes.

### Multi-omics integration highlights the transfer and the activity of bacteria from the oral cavity to the gut

Since a lower abundance of *S. salivarius* was found in both the oral cavity and the gut, we sought to explore the transmission between both extremities of the gastrointestinal tract and assess the levels of transfer in our cohort. To do so, we identified and followed genomic variations with read support from both the oral cavity and the gut (see Methods). In contrast to a previous study that only looked at the transmission using MG-based strain-variants^41^, we additionally took advantage of the MT and/or MP data to identify not only transferred but also functionally active strain-variants. Furthermore, while MG and MT analyses are based on sequencing, metaproteomics provides an independent layer of information based on peptides and mass-spectrometry analyses. This provides the opportunity to strongly validate identified transferred missense variants by identifying the translated protein with the variant amino acid sequence. Using first all genomic variants (synonymous and nonsense) with read support from both the oral cavity and the gut, we identified the genera *Prevotella* and *Bacteroides* to be transferred and active at the MT level in the gut in the majority of our cohort (**Figure 5A**). The genus *Prevotella* is relatively common and abundant in the oral cavity but less prevalent in the gut. Finding it to be transferred and active is thus not surprising. In contrast, while the genus *Bacteroides* is strongly abundant in the gut, it is rarely identified in the oral cavity^9,106^. Indeed, in our study, the signal observed from *Bacteroides* mostly came from a few particular individuals and was not representative of the entire cohort.

**Figure 5.**
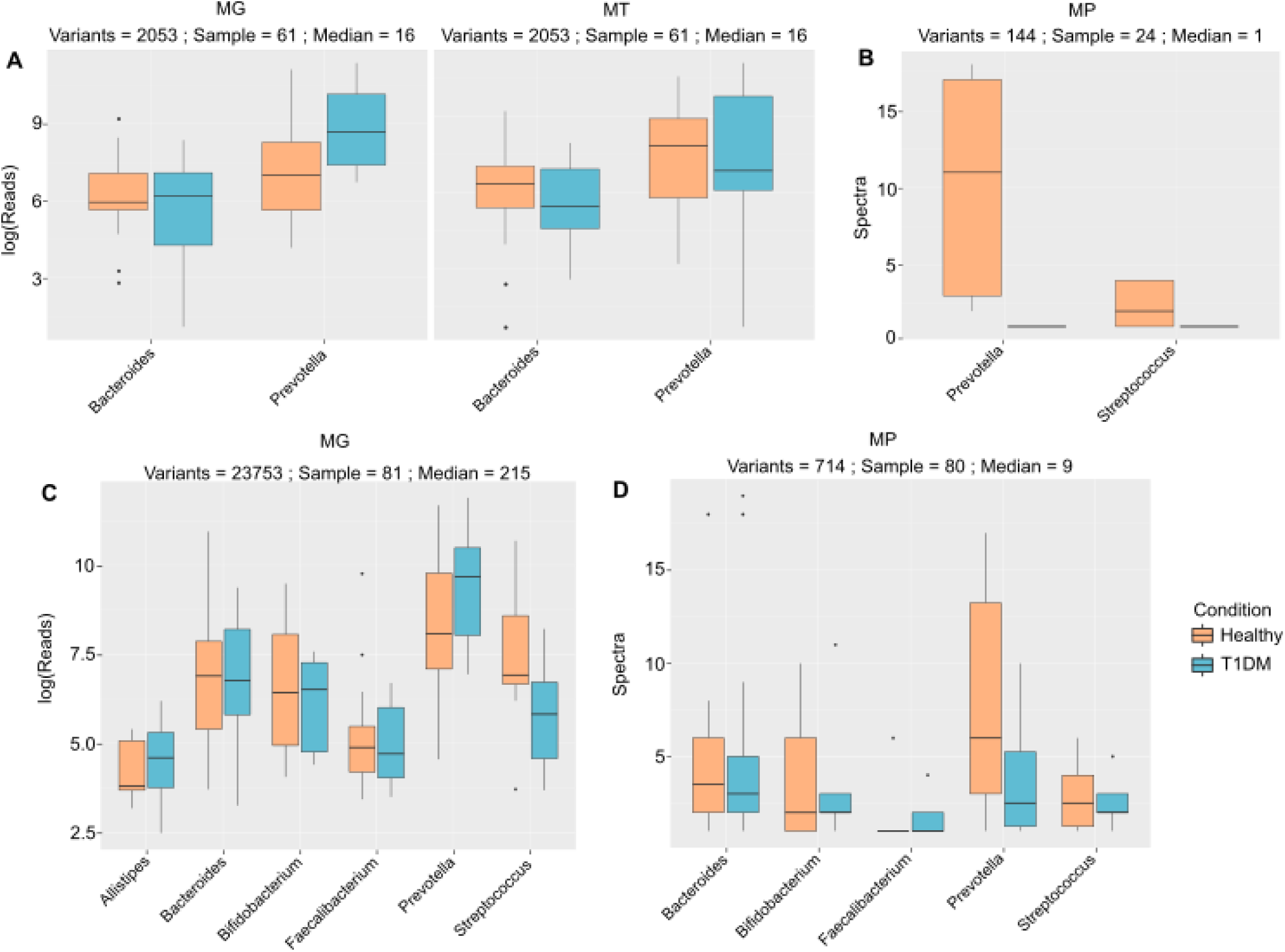
Identified variants of genera across multiple omes. The figure indicates the distribution of reads for metagenomic (MG) and metatranscriptomic (MT) abundance, and spectra for metaproteomic (MP) abundance for each set of variants associated with a taxa. The numbers on top of each box indicate the number of identified variants, the number of samples in which variants have been identified and the median number of variants per sample. Panels **A** and **B** correspond to the MG-MT supported variants while panels **C** and **D** show the MG-only supported variants. Comparisons of distributions were also performed and are represented by a light orange (healthy controls) and a light blue box (T1DM patients).

Remarkably, we identified several peptides supporting strain-variants at the MP level (**Figure 5B**), showing that we could follow, and thus validate, variants across all three omic layers. Whilst the number of variant-supporting peptides is relatively low (due notably to the typical lower depth of MP or expected lower abundance of variant peptides), their identification confirms that the taxa we find to be transferred from the oral cavity are also active in the gut. Strain-variants belonging to the genus *Bacteroides* is not identified anymore at the variant peptide level, which can be explained by the low number of samples in which *Bacteroides* was identified. More surprising is the absence of the *Streptococcus* genus using MT-supported variants but its presence at the MP level. This indicates that the representation of strain-variants belonging to the genus *Streptococcus* was too low at the MT but not at the MP level to be detected over their respective threshold (see Methods and Supplementary Table 1). Additionally, *Streptococci* are known to inhabit the upper part (small intestine) of the gut rather than the lower part (colon)^107,108^. As RNA transcripts are less stable than proteins, it is not surprising that only peptides are identifiable from taxa active in the upper gut. We thus hypothesise that the applied strict MT read abundance threshold might be too stringent to identify transferred bacteria active in the upper part of the gut and that MP support would be more appropriate. To test this, we used missense variants with only MG read support and performed the metaproteomic search including the new protein variants. We distinguish variants supported by MG from the oral cavity and MG and MT from the gut (referred to as MG-MT supported variants) and variants supported only by MG from the oral cavity and MG from the gut (referred to as MG-only supported variants). Both types of variants can be further supported at the MP level (**Figures 5B and D**).

By applying only the MG support criterion, around ten times more variants across 81 samples were found and additional genera including *Alistipes, Bifidobacterium* and *Faecalibacterium* were identified as transferred. With the exception of *Faecalibacterium*, all those taxa are commonly found at both body sites^9^. As hypothesised, strain-variants belonging to the genus *Streptococcus* were now found at the MG level (**Figure 5C**). Adding the MP layer notably confirmed the presence and the activity of the *Streptococcus* strain-variants while those from the genera *Alistipes and Faecalibacterium* (initially not found by MG-MT variants) are not found (**Figure 5D**). Metaproteomics thus essentially supports and validates the variants detected via the others omes, either due to the higher stability of proteins or to metaproteomics’ different and independent technology (e.g., it does not suffer from sequencing errors). Furthermore, as proteins are immunogenic, using metaproteomics to detect strain-variant peptides adds a valuable layer of information as proteins from the oral cavity may fuel inflammation in the large intestine.

### Streptococcus is less transmitted in T1DM in comparison to healthy controls

Being able to identify and follow variants across all omic layers and both body sites allowed us to assess the level of transfer of the different identified taxa. *Streptococcus salivarius* was found to be less abundant and less active in both the oral cavity and the gut in TIDM. While the difference is not significant, a similar trend can be observed at the transfer level for the *Streptococcus* genus. Not only does *Streptococcus* seem less transferred at the MG-only level (**Figure 5C)**, this trend seems to be further supported by lower amount of peptides, and thus a lower activity, associated to *Streptococcus* at the metaproteomic level using both the MG-MT supported variants and the MG-only supported variants (**Figures 5B and D)**. This reinforces our findings that the lower abundance of *S. salivarius* in the oral cavity and in the gut are indeed connected.

### Transmission levels strongly correlate with taxa abundances in the gut but not in the oral cavity

Correlation analyses between the MG and MT levels of the transferred bacteria and their abundance in the oral cavity and in the gut were performed in order to verify if the taxa abundances at both extremities of the gastrointestinal tract were associated. Strong positive correlations (r_s_= 0.6-0.7 at p-value <0.05) were found between the abundance (MG and MG_Only) of the transferred bacteria and their abundance in the gut, which indicates that the levels of transfer indeed influences the final abundance of the taxa in the gut (**Figure 6**). The activities (MT) were also positively correlated but at lower values (r_s_=0.4-0.5 at p-value <0.05). Interestingly, no correlations were found between the oral MG abundance of the taxa and their level of transfer (**Suppl. Figure 4**), which is consistent with the correlations found in our previous study^41^. This would suggest that the transfer rate does not simply depend on the original abundance of the taxa in the oral cavity, but rather is driven by other parameters. For example, the host physiology of the oral cavity (saliva flow-rate, glucose concentration, pH) might affect the levels of transmission along the gastrointestinal tract as well as the microbial physiology (*e*.*g*. low pH and bile acids tolerance). We therefore looked at correlations between the level of transfer and the available metadata but no significant correlations were found **(Suppl. Figure 5A and B**).

**Figure 6.**
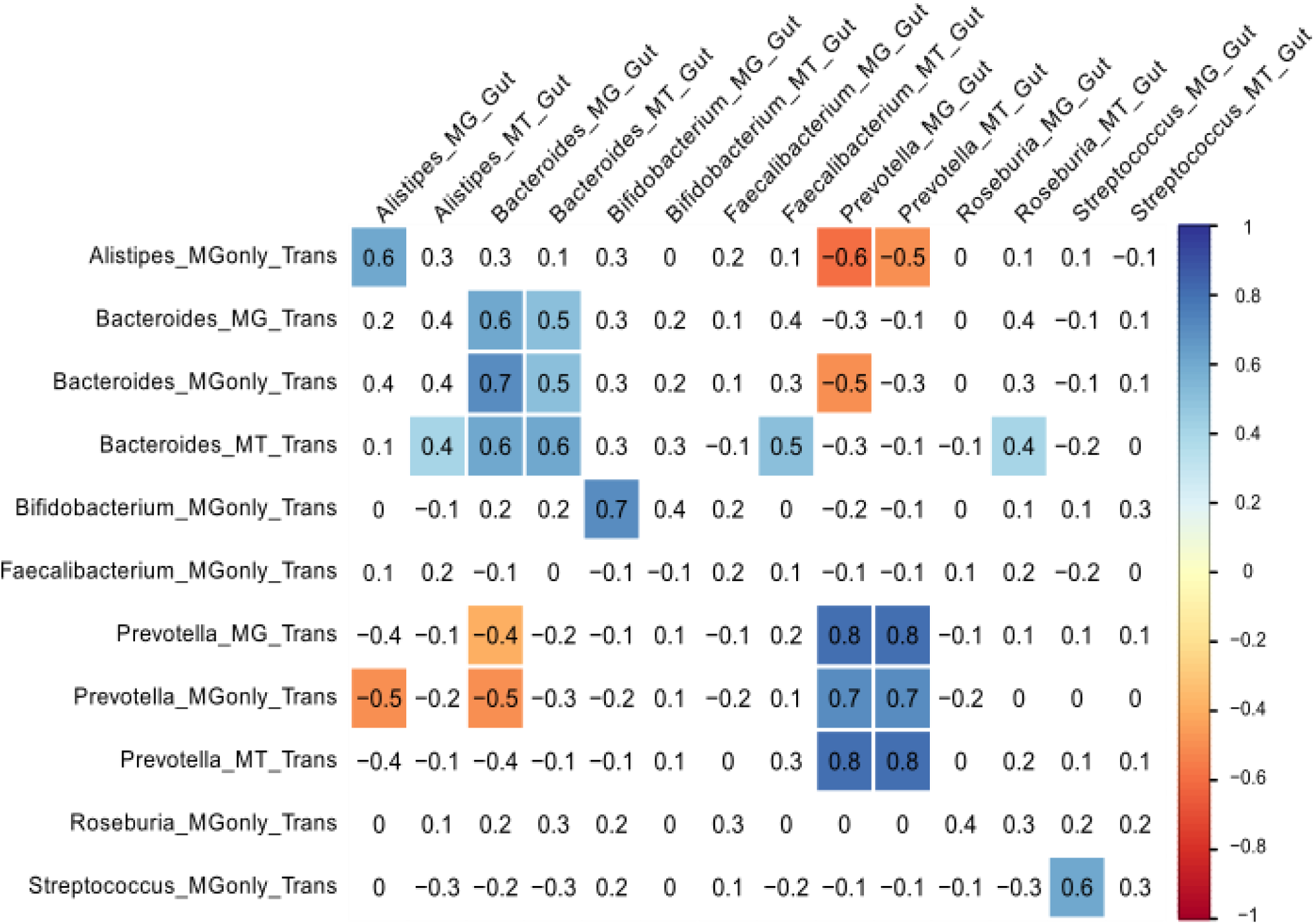
Correlations between the abundances of transferred taxa in comparison to the abundance in the gut. The figure shows the correlation between the transfer and the gut abundances. Abundances of taxa with either MG and MT labels correspond to the abundances of supported variants at the metagenomic and metatranscriptomic levels. MGonly is used if variants were supported with MG reads only and not on the MT level. Colored values indicate positive (blue) or negative (red) significant correlations (adj. p-value < 0.05). Values with white background indicate non-significant correlations.

## Conclusions

In this study, we looked at the microbiota of two important body sites at both extremes of the gastrointestinal tract, the oral cavity and the gut, and identified differences in composition, function, and transfer of bacterial taxa in a case study of familial T1DM.

In the oral cavity of T1DM patients, the abundances of different taxa strongly resembled an acidified cavity. Notably, we found a lower abundance and activity of the commensal acid-intolerant *S. salivarius* and an higher activity of the acid-tolerant pathogenic *S. mutans*, which additionally correlated with the expression of a bacteriocin, highlighting competition between the two *Streptococci* species (**Figure 2**).

In the gut, we observed lower abundance of *S. salivarius* and higher abundance of *E. coli* as well as an overall increased expression of genes involved in bacterial virulence and oxidative stress response related to the *Enterobacteriaceae* family (**Figure 3**). Besides the increased abundance and activity of *Enterobacteria*, we found further evidence of gut inflammation in T1DM through the overexpression of several human proteins involved either in the host immune response or inflammation (**Figure 4** ; **Supplementary Figure 3**).

The multi-omic data for both body sites enabled us for the first time to trace the variants and taxa across all three omic layers and thus to identify specific taxa that were both transmitted along the gastrointestinal tract, and active in the gut. This strengthened the identification of transmitted variants and brought additional evidence on actual gut colonisation by oral bacteria. We found multiple genera to be transmitted and we have highlighted the importance of using functional omic support to identify taxa active in the gut (**Figure 5**). We also discussed the limitations inherent to metatranscriptomics and highlighted how metaproteomics can be advantageously used to validate identified variants and explore the upper part of the gut.

By contextualising the information concerning oral to gut transfer in T1DM, we notably found a trend of lower levels of transmission of *Streptococcus* in T1DM patients, thereby reinforcing the notion that the lower abundance of *S. salivarius* in the oral cavity and the gut are indeed connected and both in relation to T1DM (**Figures 5B** and **D)**. However, correlations between the levels of transmission of taxa and their abundance at both body sites showed strong correlations with the gut but not with the oral cavity (**Figure 6** and **Suppl. Figure 4**). As the physiology of the oral cavity is altered in T1DM patients^42–44^ we would hypothesise that some of those factors (*e*.*g*. saliva flow-rate, glucose concentration, pH) might have a stronger influence on the transmission rate of oral microbes along the gastrointestinal tract than just their initial abundances. A follow-up study could combine different metadata measurements of the oral cavity together with the newly developed strain-variant methodology and assess if any physiological parameter influences the abundance of particular variants and their transmission rate along the gastrointestinal tract.

## Supporting information

Supplementary figues

## Data and Code availability

Metagenomic and metatranscriptomic sequencing reads can be accessed from NCBI BioProject [PRJNA289586](https://www.ncbi.nlm.nih.gov/bioproject/289586). All mass spectrometry proteomics data and results were deposited to the ProteomeXchange Consortium (http://proteomecentral.proteomexchange.org) via the PRIDE^109^ partner repository with the data set identifier PXD031579.

