## Supplementary figues for "Alterations of oral microbiota and impact on the gut microbiome in type 1 diabetes mellitus revealed by multi-omic analysis"

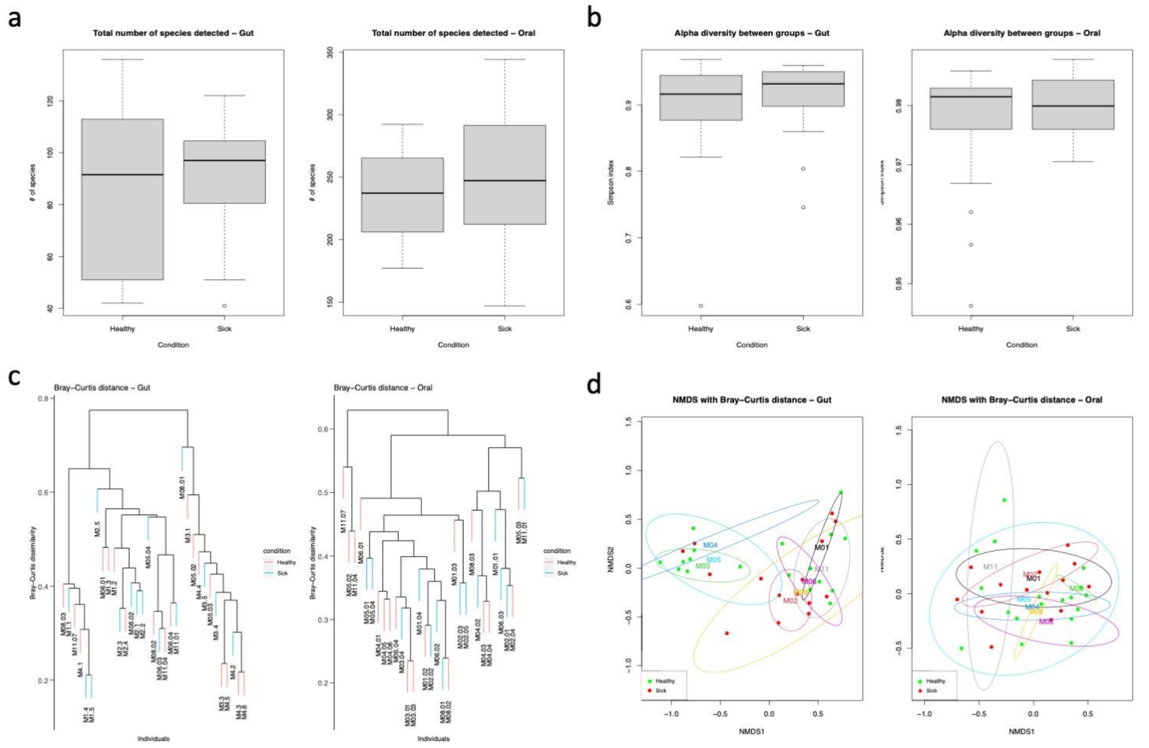


**Supplementary Figure 1. Oral and gut community structure analysis.** Box plots of species richness and alpha diversity between controls and T1DM patients (a, b). Hierarchical clustering based on Bray-Curtis distance (c). NMDS ordination based on Bray-Curtis distance (d). All analyses are based on metagenomic data.

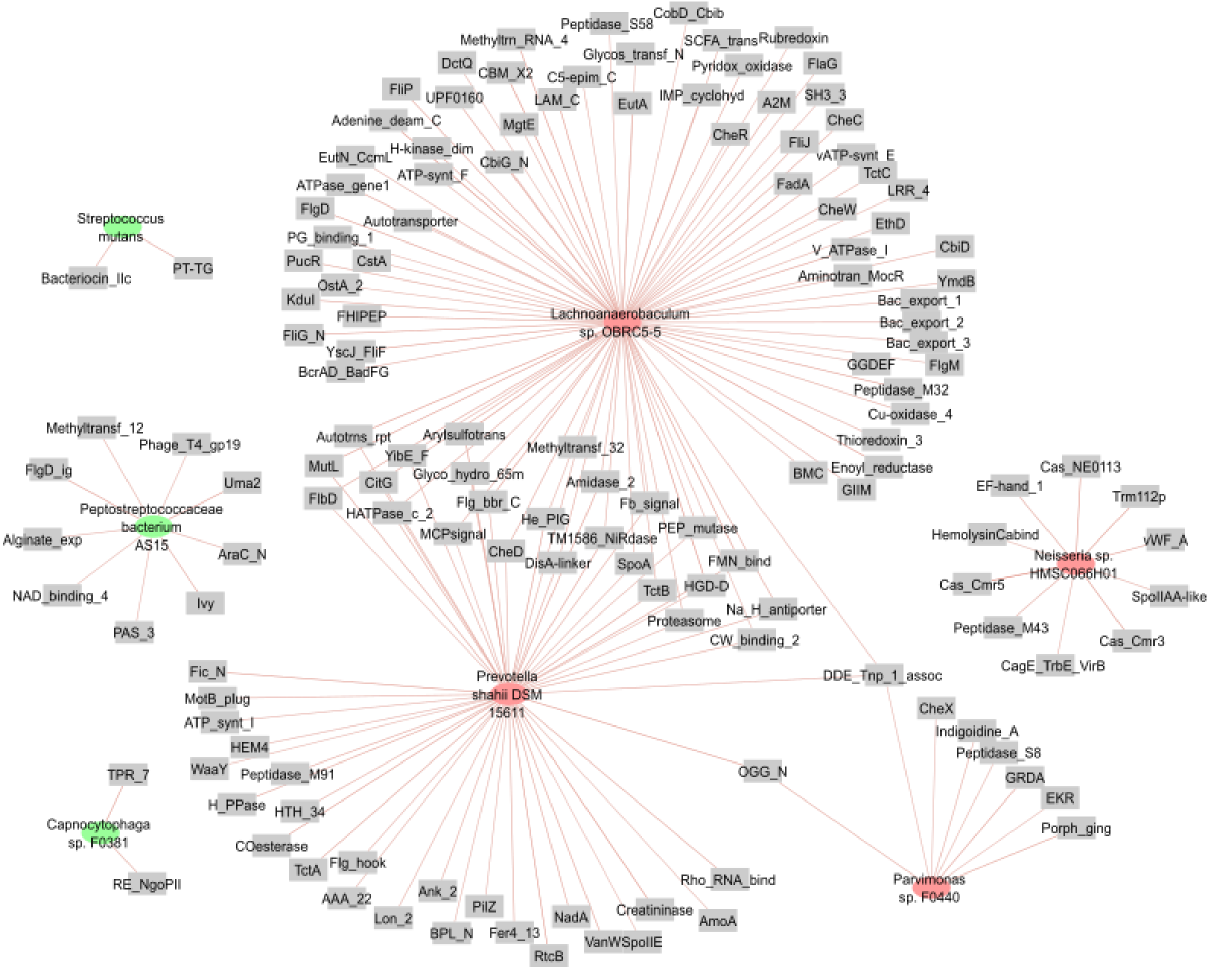


**Supplementary Figure 2. Correlation network of gene transcripts and differentially active taxa in the oral cavity.** The figure corresponds to a subset of the correlation analysis where only the differentially abundant taxa and correlation over 0.7 are plotted. Green and red nodes indicate if the taxon is up- or down-regulated in the oral cavity. Annotations are based on the Pfam database.

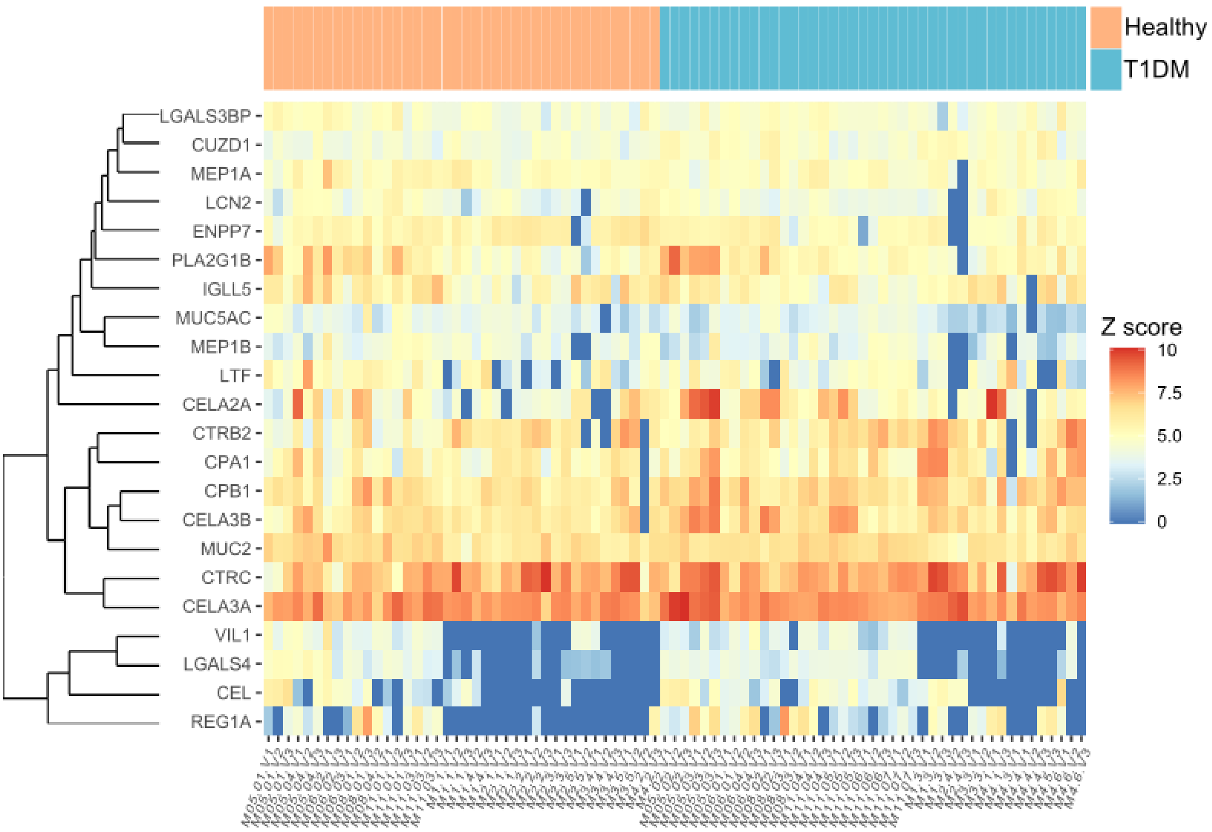


**Supplementary Figure 3. Metaproteomic differences in T1DM at the visit level.** Heatmap displaying the relative abundances of human proteins at the visit level with the highest significance in a differential analysis of T1DM. Individuals with T1DM have a purple box in the upper line on top of the columns.

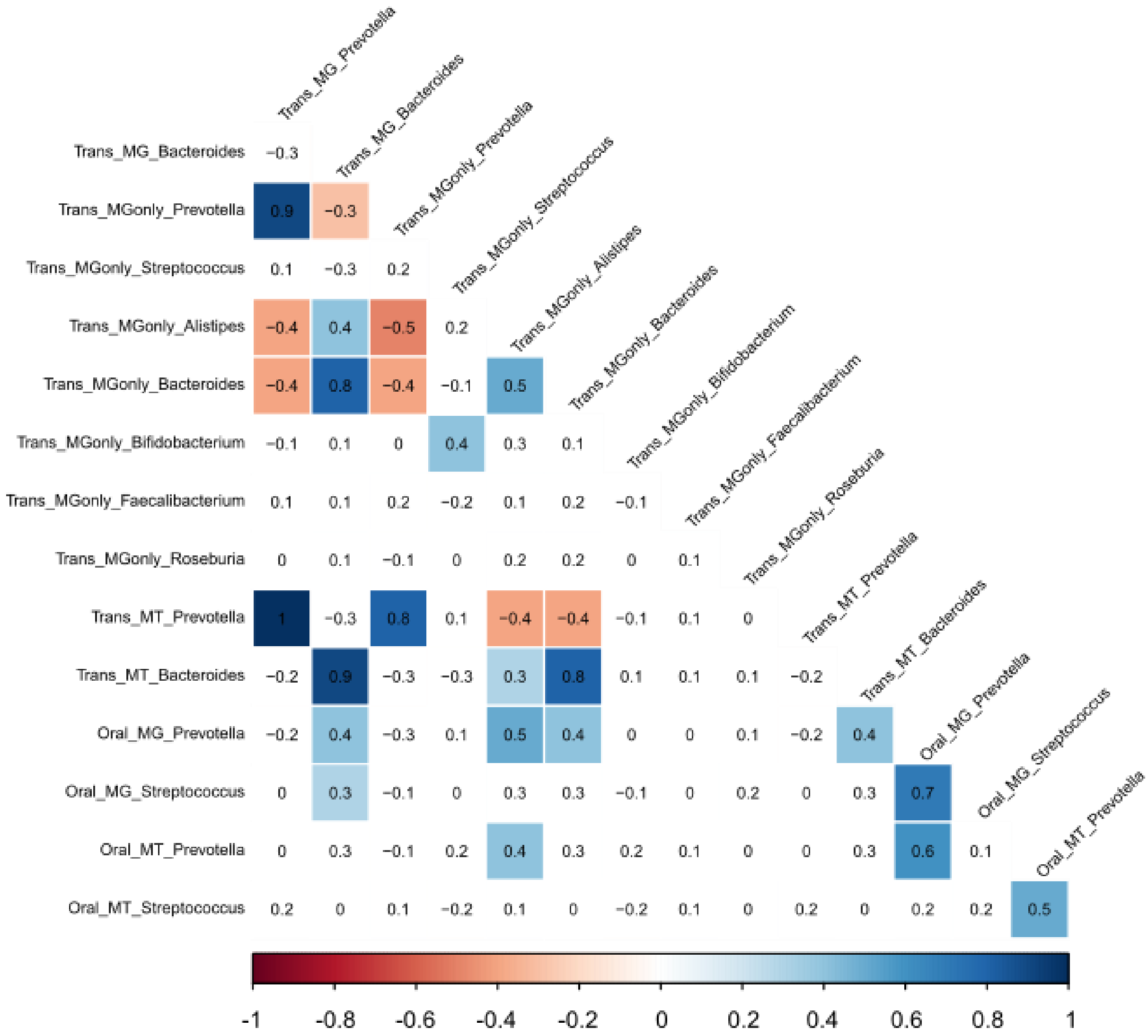


**Supplementary Figure 4. Correlations between the abundances of transferred taxa in comparison to the abundance in the oral cavity.** The figure shows the correlation between the transfer and the oral cavity. The labels with MG and MT correspond to the abundances at the metagenomic and metatranscriptomic levels for the MG-MT supported variants. MG_only is used for the MG abundances if variants supported with MG reads only and not on the MT level. Colored squares indicate a positive (blue) or negative (red) significant correlations (p-val < 0.05). White squares indicate non-significant correlations.

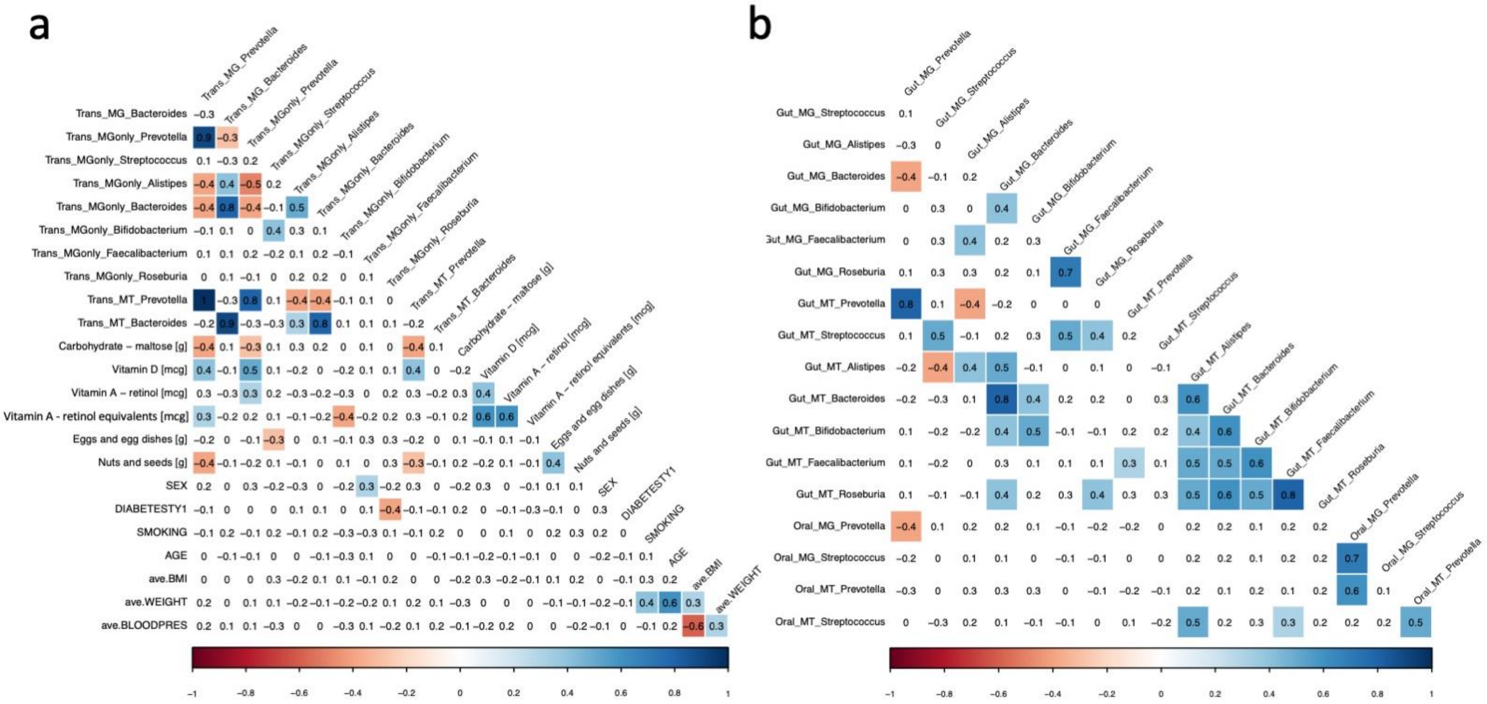


**Supplementary Figure 5. Correlation analyses of transferred genera.** Spearman correlations based on read count data of: a) Transferred genera among each other and with metadata; b) Gut with oral genera. MG: Metagenomics data, MT: Metatranscriptomics data, MGonly: Filtered on MG minimum read counts only. For dichotomous categorical data male was set to 1, female 0 for SEX, sick to 1, healthy to 0 for DIABETESTY1, and smoking to 1, non-smoking to 0 for SMOKING. Colored squares indicate a positive (blue) or negative (red) significant correlations (p-val < 0.05). White squares indicate non-significant correlations.
